# Multiple retinoic acid pathway factors function together during development of a mollusc

**DOI:** 10.64898/2025.12.01.691671

**Authors:** T. Kim Dao, J. David Lambert

**Author notes:** Department of Chemical and Biological Engineering, Princeton University, Princeton NJ, United States.

## Abstract

In developing chordate embryos, the retinoic acid (RA) pathway is involved in many key developmental patterning systems. Recently, it has become clear that at least some components of the RA pathway are more ancient than chordates. However, the participation of these components in an RA pathway, and the role of such a pathway in developing non-chordate embryos remain unclear. Here, we demonstrate the presence of an extensive set of RA pathway components in the genome of the mollusc *Tritia obsoleta,* and examine their expression using *in situ* hybridization. We then examined the function of multiple RA pathway genes using RA treatments, drug treatments and morpholino (MO) knockdowns. These manipulations impacted a similar set of structures in development, especially the shell and the digestive tract, indicating that the retinoic acid pathway is functional in *Tritia.* Together, this is the most comprehensive evidence yet for an RA signaling pathway functioning in embryogenesis of a non-chordate.

## INTRODUCTION

Retinoic acid (RA) is a lipophilic bioactive derivative of Vitamin A that plays many important roles in vertebrate development and physiology. During embryogenesis, RA forms concentration gradients and acts as a morphogen (reviewed in Cunningham and Duester, 2015). This occurs in the hindbrain, where RA concentrations help delineate and specify rhombomeres (Dupe and Lumsden, 2001; Lee and Skromne, 2014; Niederreither and Dolle, 2008), in the presomitic mesoderm, where an RA gradient helps to position the determination front in somitogenesis (Duester, 2007), and in limb development, where RA gradients exist across the distal limb in the anterior-posterior axis, and along the proximo-distal axis (Giguere et al., 1989). In some of these contexts, RA acts through Hox genes to confer patterning information (Kashyap et al., 2011; Kashyap et al., 2013; Langston et al., 1997). In the development of the nervous system, RA is required for specification of particular cell types, like motor neurons, and is also involved in coordinating cortical differentiation, neural outgrowth and axon guidance (Corcoran and Maden, 1999; Harrison-Uy et al., 2013; Janesick et al., 2015; Niederreither and Dolle, 2008; Novitch et al., 2003). RA is also necessary for normal development of multiple other organs in vertebrates, including the pancreas, foregut, kidney, heart and eye (e.g. Mark et al., 2006).

The molecular machinery for synthesis, transport, transcriptional activity and degradation of RA has been well-characterized in vertebrates (reviewed in Ghyselinck and Duester, 2019). Figure 1A shows the core RA pathway components that we address in this report. Vertebrates acquire vitamin A via their diets mainly in the form of β-carotene. This molecule is then broken down into retinal by β-carotene-15,15′-monooxygenase 1 (BCMO1). In RA signaling, extracellular retinol binding proteins like RBP4 bind to retinol and transport it to the target cells. Since retinol, retinal and retinoic acid are poorly soluble in aqueous environments, cellular binding proteins such as cellular RA-binding protein (CRABP) chaperon them inside a cell. RA is synthesized from retinol via 2 oxidative steps. The first step is the reversible conversion of retinol into retinal by retinol dehydrogenases (RDHs). In mouse, this is the rate-limiting step in embryonic RA synthesis (Sandell et al., 2007). In the second step, retinal is irreversibly converted into RA by retinaldehyde dehydrogenases (RALDHs). RA then binds to the RAR-RXR heterodimers. This ligand-receptor complex bind to the retinoic acid response elements (RAREs) upstream of genes and either activate or suppress gene transcription. Cytochrome P450 26 (CYP26) proteins degrade RA.

**Figure 1.**
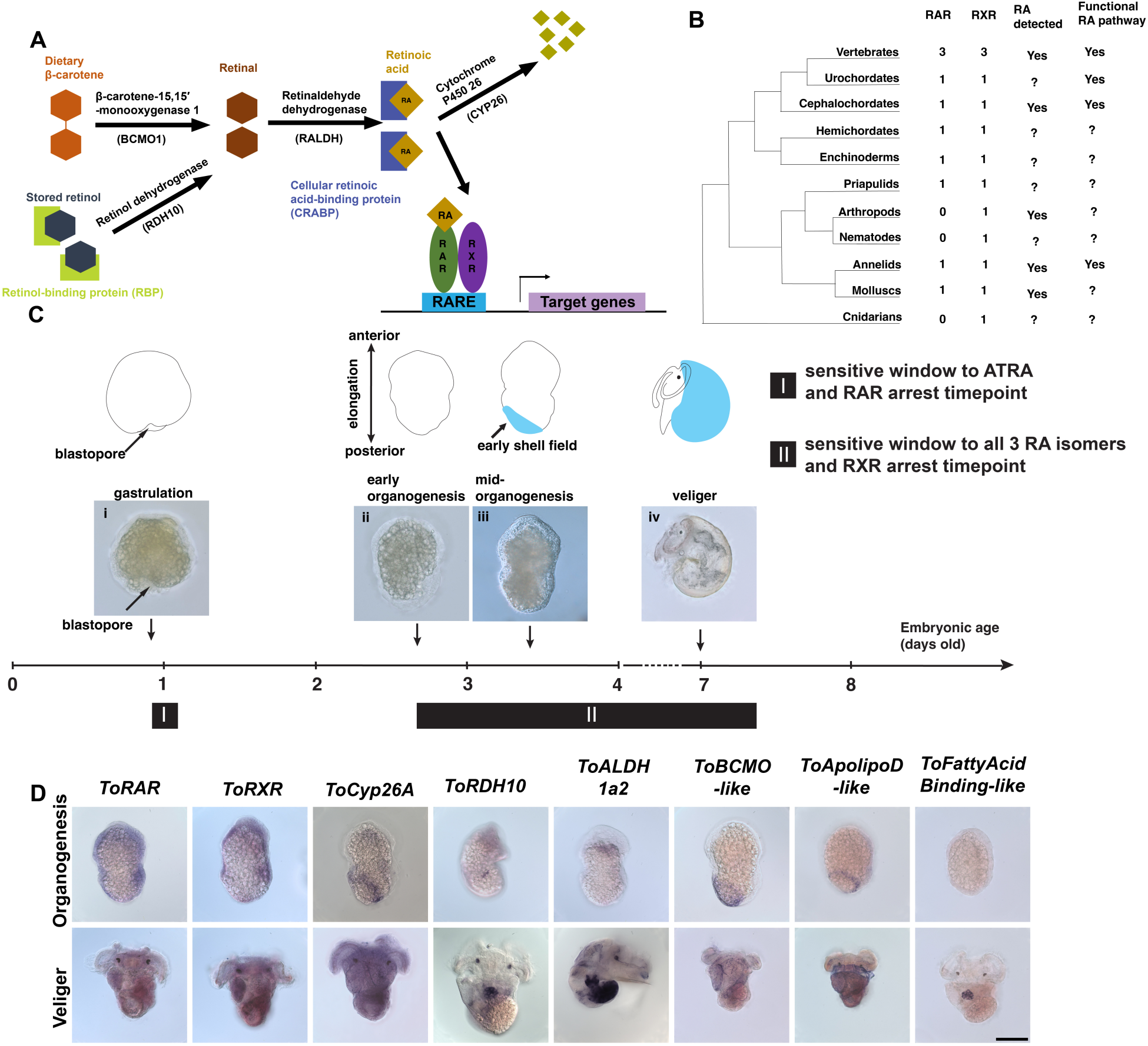
(A) A simplified model of the vertebrate RA signaling pathway. (B) Summary of *RAR*, *RXR*, RA and RA pathway characterization across the animal kingdom. (C) Embryonic and shell development of *Tritia obsoleta* in relation to retinoic acid sensitivity. (i) Wildtype late gastrulation embryo, around 24 hours after embryos are laid. (ii) Wildtype early organogenesis embryo, about 2.8 days after egg laying. The shell is one of the first organs to form during organogenesis. (iii) Wildtype mid-organogenesis embryo, around the 3.5-day-old mark. The early shell field has formed on the left side of the embryo. (iv) Wildtype veliger larvae, around 7-days after laying. Once the veliger has depleted its yolk it has a functional digestive system and hatches into the water column. (D) Expression of putative RA signaling pathway components during organogenesis and veliger stages. Scale bar: 100 μm

RA exists in several isomers in vertebrates. All-trans retinoic acid (ATRA) is the dominant isomer in vertebrates, and binds to and activates the RARs. 9-cis RA has activity in some contexts and has high affinity for RXR; however, detection of 9-cis in vivo in mammals has been elusive (Kane et al., 2005; Pappas et al., 1993; Schmidt et al., 2003). 13-cis RA also has activity in some situations (Tang and Russell, 1990; Zouboulis et al., 1991).

Until relatively recently, retinoic acid signaling was thought to be a chordate innovation. The two most powerful invertebrate model organisms, the fruit fly and the soil nematode do not have retinoic acid signaling, and evidence for RA function in other invertebrates was sparse and based on exposure to exogenous RA, which does not necessarily reflect endogenous function (Duester, 2017; Gutierrez-Mazariegos et al., 2014b; Kastner et al., 1995). So, it was surprising, as more and more invertebrate genomes were sequenced, to find that most animal groups do in fact have orthologs of many of the genes involved in RA signaling in vertebrates, including RAR, RXR, Cyp26, RALDHs and RDH/SDR genes (Albalat and Canestro, 2009; Canestro et al., 2006; Gutierrez-Mazariegos et al., 2014b; Sobreira et al., 2011). Retinoic acid has been detected in vertebrates, cephalochordates, arthropods (*Locusta migratoria*), annelids (*Platynereis dumerili*), and molluscs (Nowickyj et al., 2008). While RA signaling pathway is extensively studied in chordates (vertebrates, urochordates and cephalochordates), little is known about the functions of the pathway in non-chordate phyla. Treatment with retinoic acid induces mitosis in cleavage-arrested apical plate cells in the gastrula stage of the gastropod mollusc *Lymnaea stagnalis* (Creton et al., 1994). Retinoic acid also has various effects on neuronal development, neurite development, axon guidance and synaptogenesis in the *Lymnaea* nervous system (e.g. Dmetrichuk et al., 2006; Farrar et al., 2009). Imposex is a well-studied phenomenon where male sex characteristics develop in female snails due to environmental organotin exposure (Bryan et al., 1988; Smith, 1981). In adult *Nucella lapillus*, 9-*cis* retinoic acid promotes imposex as efficiently as organotin (Castro et al., 2007), and RA has been implicated in normal sex organ development in *Crepidula fornicata,* another gastropod mollusc (Lesoway and Henry, 2021). The most detailed functional characterization of the RA pathway in development outside chordates was performed in the annelid *Platynereis dumerili.* In this embryo, retinoic acid seems to play a role in neurogenesis and axon growth in the medial neuroectodermal tissues, and they confirmed the functional requirement for *RAR* and *RXR* (Handberg-Thorsager et al., 2018).

Despite these observations, it remains unclear how RA signaling functions in non-chordates, and thus how the pathway evolved in the vertebrate lineage. This is mainly because few functional studies of the pathway have been published in non-chordate systems. Those that have been performed have been limited to perturbation of the receptors; the other pathway components have not been functionally assessed. To understand the evolution of the retinoic acid pathway, we need to characterize the functions of the retinoic acid pathway in additional phyla outside chordates and define the set of genes that participate in the pathway in non-chordates.

In this study we use the model mollusc embryo of *Tritia obsoleta* to examine the expression of a set of putative RA pathway genes. We then take advantage of the strengths of the *Tritia* embryo for functional analysis of embryonic development (e.g. Gharbiah et al., 2009) to test the effects of specific gene knockdowns of four core RA pathway genes, alongside treatments with exogenous RA, and an RXR inhibitor. We find that most treatments cause a very similar spectrum of defects that are centered on the development of the shell. Together, similarities between these phenotypes, and overlap in the expression patterns in the shell and dorsal mantle cavity, provide the best functional evidence to date of an extensive RA pathway in a non-chordate embryo.

## RESULTS

### Putative RA pathway components in Tritia

We detected clear orthologs of the main actors of the RA pathway and RA transportation genes in the *Tritia* transcriptome using the reciprocal best BLAST hit approach (Wolf and Koonin, 2012; summarized in Table 1). *ToRAR*, *ToRXR*, *ToCYP26A*, *ToRDH10* and *ToALDH1a2* were reciprocal best hits to their respective orthologs. These genes have been identified in most other non-chordates where they have been sought, indicating that they were present in the bilaterian ancestor (Albalat, 2009; Albalat and Canestro, 2009; Campo-Paysaa et al., 2008; Shea et al., 2004).

**Table 1:** Putative *Tritia* RA pathway genes reported here.

| Gene name | BLAST-based “reciprocal best hits” results | *In situ* expression patterns |
| --- | --- | --- |
| *ToRAR* | Reciprocal best hit with mouse ortholog | ubiquitous |
| *ToRXR* | Reciprocal best hit with mouse ortholog | ubiquitous |
| *ToCYP26A* | Reciprocal best hit with mouse ortholog | shell field |
| *ToRDH10* | Reciprocal best hit with mouse ortholog | cluster of cells in dorsal midline of mantle cavity |
| *ToALDH1a2* | Reciprocal best hit with mouse ortholog | digestive glands |
| *ToBCMO1-like* | Best hit to members of the BCMO1, BCDO2, Rpe65 paralogy group | shell field |
| *ToApolipoD-like (RBP4-related)* | Best hit to “ApolipoproteinD”, member of vertebrate paralogous protein family including Retinol Binding Protein 4 | shell field |
| *ToFattyAcidBinding-like (CRABP-related)* | Best hit to “Fatty Acid Binding Protein” in a protein family that is related to vertebrate CRABP | cluster of cells in dorsal midline of mantle cavity |

We also identified another class of putative RA pathway proteins that have received less attention in non-chordate groups. These are cases where the vertebrate RA-specific protein appears to be the result of duplications in the vertebrate lineage, so invertebrates do not appear to have the specific paralog that has been characterized in vertebrates, but we can find proteins that appear to be orthologous with one of the vertebrate paralogs of the gene that has been implicated in RA function. For these proteins the closest BLAST hit for the *Tritia* sequence is one of multiple mammalian paralogs. In these cases, our working hypothesis is that the ancestral gene may have participated in RA signaling, but that in the vertebrate lineage, there was a sub-functionalization, so a new duplicate was further specialized in its RA-related function. For instance, it has already been argued that vertebrate *RBP4* evolved from an apolipoprotein that is conserved across animals (Diez-Hermano et al., 2021; Sanchez et al., 2003) and the closest *Tritia* sequence to *RBP4* is an ortholog of this apolipoprotein in mice, which is paralogous with *RBP4*. So, if there was a functional RA pathway in non-chordates, it seems possible that this ApoD-like lipocalin may have served a similar role to RBP4 in that ancestor, and after the locus was duplicated twice with the rest of the genome at the base of the vertebrates, the presumptive RBP4 duplicate evolved to specialize in RA transport. In this scenario, we would argue that a best hit to any of the vertebrate paralogs is consistent with orthology to RBP4, and makes the gene a candidate RA pathway participant. *ToBCMO1-like* has highest similarity to vertebrate BCMO1 genes, but is also very similar to other vertebrate paralogs of BCMO1, namely Rpe65 and BCDO2 (Albalat, 2009); thus, we suspect that it is the ortholog of the gene that duplicated to create these paralogs in vertebrates, and may have a role in RA signaling in *Tritia.* Similarly, vertebrate cellular retinoic acid-binding proteins (CRABPs) belong to a large group of intracellular lipid-binding proteins (iLBPs) (Schaap et al., 2002). Putative iLBPs have been proposed in protostomes to be potentially RA-binding, even if they are not directly orthologous to vertebrate CRABPs (Folli et al., 2005; Gu et al., 2002; Mansfield et al., 1998; Wang et al., 2007). A gene we named *ToFattyAcidBinding-like* in the *Tritia* transcriptome was a best hit to a fatty acid binding protein in mouse that is related to CRABP1.

### Expression patterns of RA pathway components and RA detection

To examine the expression patterns of these putative components of the retinoic acid pathway, we performed mRNA *in situ* hybridization for *ToRAR*, *ToRXR*, *ToCYP26A*, *ToBCMO1*, *ToRDH10* and *ToALDH1a2* at two developmental stages, organogenesis (3.5 - 4 day-old) and veliger (7 day-old) (Figure 1D). The heterodimeric receptors *ToRAR* and *ToRXR* were expressed ubiquitously at both stages. *ToCYP26A* and *ToBCMO1* mRNAs were specifically expressed on the left-posterior side of the body in the shell field at organogenesis (3.5 - 4-day old) stage. Both transcripts lost their specificity and became ubiquitous at the veliger stage. *ToRDH10* mRNAs started from a cluster of cells on the right side of the body at the start of organogenesis, and as the embryos matured into veliger stage, *ToRDH10* became specifically expressed in a cluster of cells in the dorsal midline of the mantle cavity on the dorsal side of the mantle cavity. *ToRALDH* was faintly expressed near the head during organogenesis and became specific to the digestive glands as embryos matured into veligers. *ToApolipoD- like* (RBP4-related) was expressed in the shell field during organogenesis stage embryos. *ToFattyAcidBinding-like (CRABP-related)* mRNA was not detected during organogenesis, but in the veliger it was in a cluster of cells on the dorsal midline in the mantle cavity. These patterns indicate that there is spatial control of RA levels in the developing *Tritia* embryo. The presence of multiple putative components in the shell and dorsal mantle cavity suggest that these structures may be foci of RA signaling.

### Elevating the RA level by RA treatment and ToCYP26A knockdown

We treated embryos with the 3 RA isomers at various developmental stages and durations of treatment. These experiments showed that were two distinct windows of sensitivity, one at the gastrulation stage, and a second that started at the elongation phase of early embryogenesis and lasted about 5 days. Treatments that did not include one or both windows resulted in wild-type larvae, and these treatment windows are the minimum amount of time that are required to generate the reported phenotypes. During gastrulation, embryos treated with ATRA showed a dramatic and highly reproducible phenotype: a yolky cellular “tail” extended from the vegetal pole of the animal and eventually broke off (Figure 2A [i] & [ii]). These tails formed quickly, usually within 15 minutes of treatment. Aside from yolky “tails”, we frequently observed large round cells that had fallen off embryos. Larva derived from these “tail”-squeezing embryos grew up to be smaller but overall wild-type (Figure 2A[iii]). The phenotype was specific only to ATRA and the gastrulation stage. We stained treated and control embryos with fluorescently labeled phalloidin to visualize the actin cytoskeleton, but did not observe any differences (not shown). To our knowledge, this effect has not been reported before, but there may be related findings in the literature. Embryonic cells of the sea urchin *Hemicentrotus pulcherrimus* developed pseudopodial cables when treated with ATRA (Kuno et al., 1999). RA treatment of neurons in various systems, including the mollusc *Lymnaea,* causes neuronal projections (Dmetrichuk et al., 2008).

**Figure 2.**
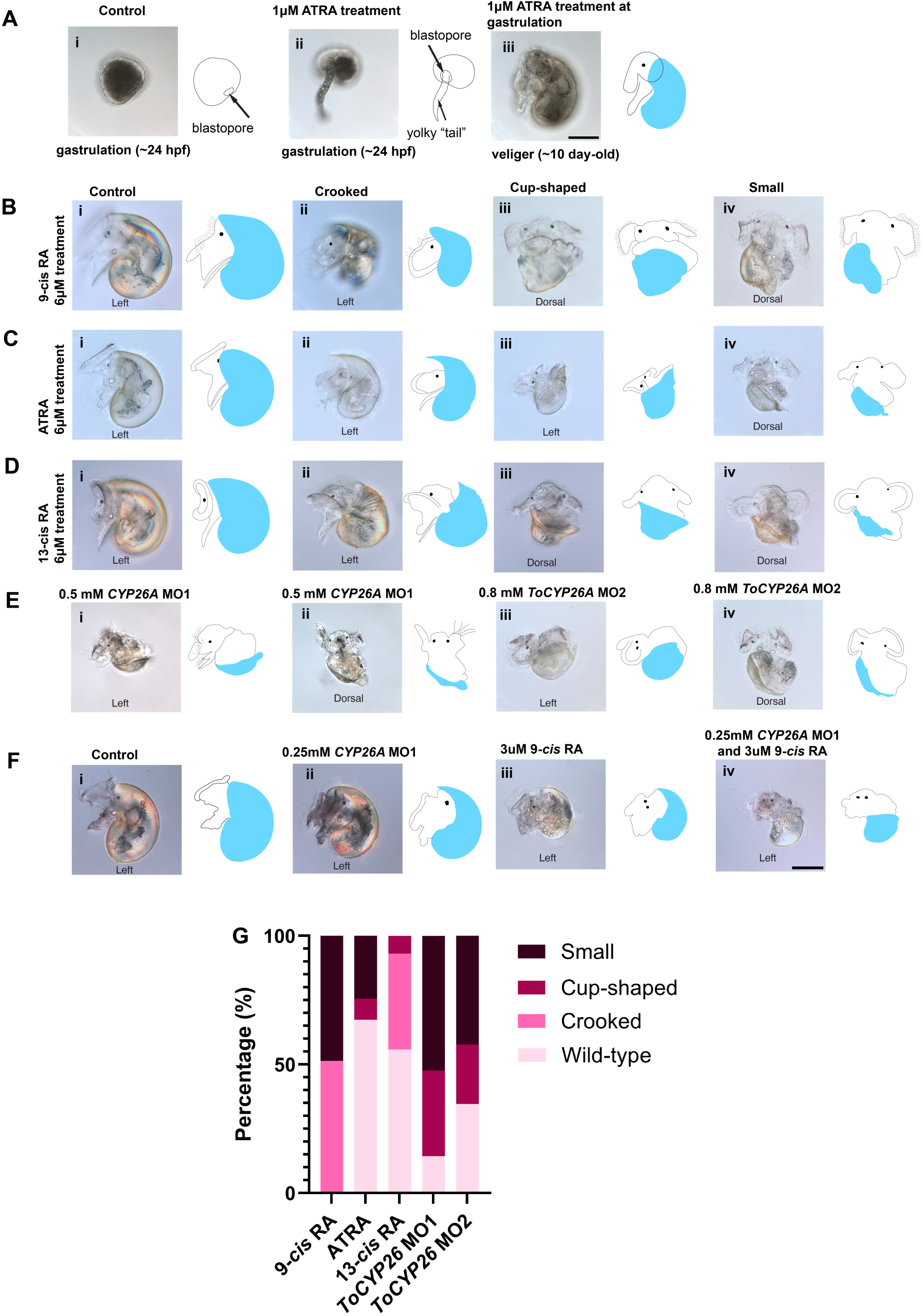
Effects of increasing retinoic acid levels by retinoic acid treatments and *ToCYP26A* knockdowns in *Tritia*. (A) Treatment with ATRA at gastrulation stage resulted in a cytoplasm “tail” that was extruded from the embryo at the blastopore (A[ii]). This was not observed in DMSO control embryos (A[i]). Embryos exhibiting the extruded tail grew up to be overall wild-type (Figure A[iii]). Scale bar: 100 μm (B, C, D) Shell defects from treatments with RA isomers. Treatments of early organogenesis embryos for 5 days with 9-*cis* RA (Figure B), 13-*cis* RA (Figure C), and ATRA (Figure D) cause shell defects (compare to DMSO control embryos (Figure B,[i], C[i], D[i]). See text for details. (Blue: shell). Scale bar: 100 μm. (E) Shell defects from *ToCYP26A* morpholino knockdown. *ToCYP26A* MO controls were wildtype, compare to controls in A, B, D (not shown). Knockdown of *ToCYP26A* with 2 non-overlapping morpholinos result in similar shell defects. Shell phenotypes in *ToCYP26A* knockdown range from wild-type, cup-shaped to small. (Blue: shell) (F) Test for synergy between 9-cis treatment and *ToCYP26A* morpholino knockdown. Embryos injected with 0.25mM *ToCYP26A* MO1 were smaller in general but their shells were normal: they coiled and covered their bodies. Shells of embryos treated with 3 μM 9-*cis* RA were significantly smaller; however, they still covered the body. Embryos injected with 0.25mM *ToCYP26A* MO1 and treated with 3 μM 9-*cis* RA were very small, uncoiled and did not cover the body. Scale bar: 100 μm (G) Summary of shell phenotypes in RA treated embryos and *ToCYP26A*MO animals, which displayed shell defects in 4 categories: wild-type, crooked, cup-shaped, and small. See text for details.

The second window of RA sensitivity started at the early organogenesis stage (∼ 2.8 - 3 days after egg laying), when embryos elongate from the round shape of the gastrula and shell formation begins. It lasted for about five days after this stage, to 7-8 days after egg laying, and caused a specific suite of phenotypes, most obviously in the shell (described below). RA treatments generally slowed development and yolk consumption. After the treatment ended, embryos were reared until they depleted their yolk, which generally corresponds to when they have finished organogenesis and developed into a hatchling veliger larva. This took 7-8 days for control embryos, and 10-11 days for RA-treated embryos.

Other windows of treatment did not permanently disrupt development. For instance, exposure from 2-4 days somewhat delayed development, but not as drastically as the 5 day window, and ultimately led to normal morphology. When the treatments continued from 2-8 days, larvae had the same defects as embryos treated from 3-8 days. Embryos treated with RA at 4 days old or later did not have shell defects, even when the treatments went on for 5 days. Thus, the phenotypes we saw required treatments that spanned early organogenesis to 8 days after egg laying. We observed this window with ATRA, 9-*cis* RA and 13-*cis* RA.

Treatment with each of the three isomers resulted in the same very specific set of phenotypes. First, the shell seemed to be malformed to varying degrees. We classified the shell morphologies into 4 categories: wildtype, crooked, cup-shaped, and small (Figure 2B). Crooked shells still grew to cover the body and were coiled, but the shell was bent, creating an angle. In some cases, crooked shells had biomineral ridges running across the shell perhaps suggesting an interruption of normal shell growth. For cup-shaped shells, the shell grew from the left to the right side of the embryo as normal, but these shells did not cover the body and did not coil. The most severe shells, which we categorized as small, stayed on the left side of the body, similar to the shell of an early organogenesis embryo. Of the three isomers, 9-*cis* RA consistently led to the most severe shell phenotype, ATRA was intermediate, and 13-cis RA treatments were least severe (Figure 2G). Besides the shell, we scored the effects of RA treatments on the other readily scorable larval structures that we routinely score (Table 2). 9-cis RA treated embryos often had defects in the digestive glands, stomach, intestine and sometimes the larval retractor muscle. ATRA and 13-cis treatments were similar but less severe. We observed less severe intestinal defects in embryos with cup-shaped shell, suggesting the defects in the intestine might be secondary to the shell defect.

**Table 2.**
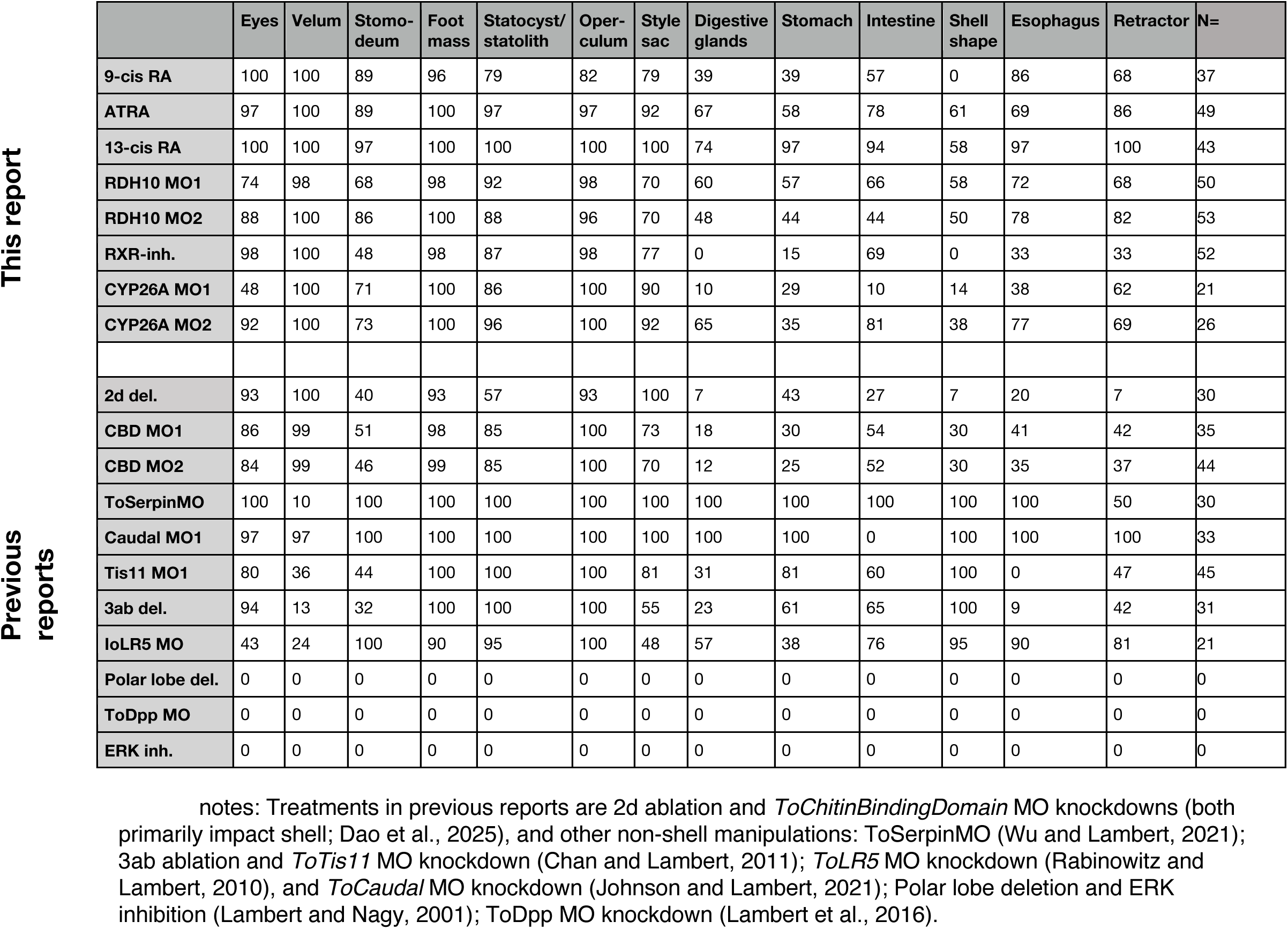
Phenotypic effects on larval organs.

Indeed, we recently reported that blocking the development of the shell, either by ablation of the 2d cell, or by knockdown of a shell-specific transcript, caused secondary defects in the digestive glands, stomach, intestine and retractor muscle (Dao et al, 2025; summarized in Table 2; 2d deletion and CBDMo1 and CBDMo2 treatments).

Given the dependence of these organs on normal shell development, these results indicate the exogenous RA treatment in this window causes consistent defects in shell development along with expected defects in these other structures.

Since *CYP26* proteins are predicted to degrade RA, knockdown should have similar effects as adding exogenous RA. We knocked down *ToCYP26A* using a translation-blocking morpholino oligo *ToCYP26A* MO1. We injected the zygote with 0.5 mM *ToCYP26A* MO1 and reared them until they had depleted their yolk and completed organogenesis (∼ 10 days, 2-3 days longer than controls). *ToCYP26A* MO1-injected embryos displayed a range of shell defects that were similar to RA treatment, i.e. small and cup-shaped shells (Figure 2E, G). They also displayed defects in the digestive glands, intestine, stomach and retractor; Table 2). To confirm that the knockdown effects of *ToCY26A* MO1 were specific, we used a second non-overlapping *ToCYP26A* morpholino, *ToCYP26A* MO2. This caused similar defects in the shell development (Figure 2E, G). *ToCYP26A* MO1 and MO2 knockdown had generally similar phenotypes but *ToCYP26A* MO1 was more severe for some organs (Fig. 2E,H and Table 2). The similarity of *ToCYP26A* MO knockdown phenotypes to RA treatment phenotypes suggests that *ToCYP26A* is participating in RA signaling in this embryo.

To further test whether *ToCYP26A* was participating in RA signaling, we looked for a synergistic effect between exogenous RA and *ToCYP26A* knockdown (Figure 2D). Embryos injected with 0.25 mM *ToCYP26A* MO1 (*vs*. 0.5 mM in 2C) were slightly smaller but otherwise normal. Embryos treated with 3 uM 9-*cis* RA (*vs*. 6μM in Figure 2B) had significantly smaller shells, but they still covered the body and were otherwise wildtype. In contrast, embryos injected with low *ToCYP26A* MO1 and treated with low RA concentration were very small. Their shells were uncoiled and did not cover the body, and they otherwise resembled 9-cis RA and *ToCYP26A* MO1 phenotypes. This result suggests that RA and *ToCYP26A* are functionally interacting.

### ToRDH10 knockdown and shell development

*RDH10* synthesizes retinal, a RA precursor. In mouse embryos, *RDH10* is the essential rate-limiting gene in RA biosynthesis (Sandell et al., 2007). *RDH10* with mutations in its catalytic domains were lethal in midgestational mouse embryos (Rhinn et al., 2011). We knocked down *ToRDH10* with a morpholino (*ToRDH10* MO1). *ToRDH10* MO1 knockdown caused a range of shell phenotypes that are very similar to those caused by RA treatments; they can also be categorized into normal, crooked, cup-shaped, and small shells (Figure 3A). They also often had defects in the other organs that were affected in 9-cis RA and *ToCYP26A* MO phenotypes (e.g. digestive glands, stomach, and intestine; Table 2). To verify the specificity of *ToRDH10* MO1, we knocked down *ToRDH10* using a second non-overlapping morpholino (*ToRDH10* MO2), which generated very similar phenotypes (Fig. 3B; Table 2).

**Figure 3.**
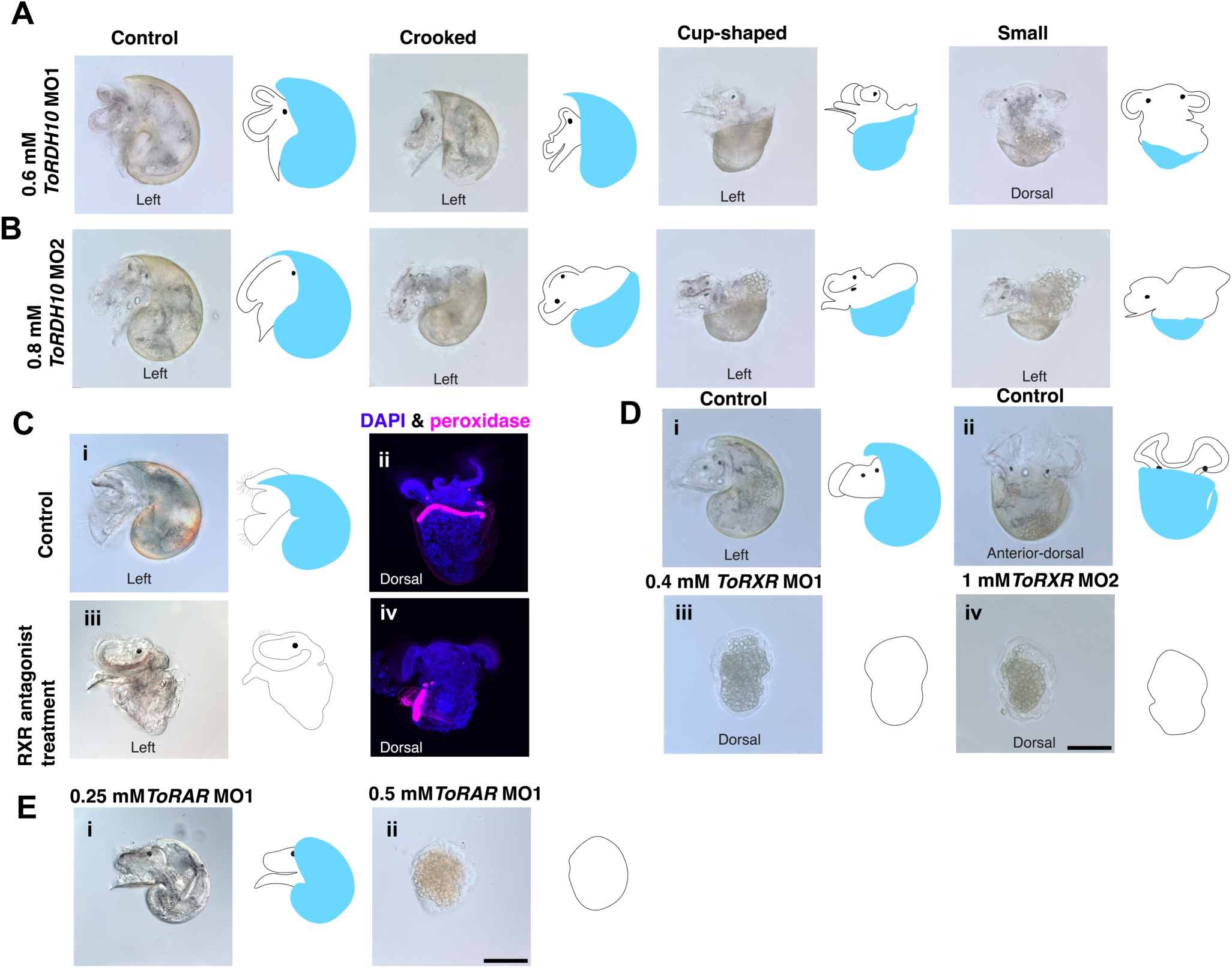
Phenotypic effects of *ToRDH10*, *ToRXR*, and *ToRAR* pertubations. (A, B) Shell defects in *ToRDH10* morpholino knockdown embryos. Knockdown of *ToRDH10* using two non-overlapping morpholinos resulted in similar shell phenotypes: crooked, cup-shaped, and small. (C) Shell and periostracal membrane defects in *ToRXR* antagonist-treated embryos. *(i)* In freshly fixed larvae, the shell shows birefringence under polarized light, observed in this control as a shiny, golden appearance. (This is not observed in other controls because minerals are lost from the shell matrix with time after fixation.) After RXR antagonist treatment no birefringence is observed. (ii and iv) Peroxidase staining of the periostracal groove in a control (ii) and RXR antagonist larvae, showing that the periostracal groove is smaller and abnormally positioned after treatment. (D) Embryonic phenotypes in *ToRXR* morpholino knockdown embryos. *ToRXR* knockdown morpholino embryos were arrested at late organogenesis stage. Scale bar: 100 μm (E) Embryonic phenotypes in *ToRAR* knockdown embryos at two different injected concentrations (0.25 mM or 0.5 mM) of *ToRAR* MO. [i] is wildtype, [ii] is arrested near the end of gastrulation. Scale bar: 100 μm

### RXR antagonist treatment and ToRXR morpholino knockdown

The functional roles of RXR in protostomes are likely to be complex. It can dimerize with NRs other than RAR, and it can exert effects with and without a ligand (e.g. RA). We first tested the effects of blocking normal ligand binding by treating organogenesis-stage embryos with UVI-3003, a pharmacological antagonist of RXR for 1-3 days. These animals had consistent defects in the shell, as well as the digestive glands and stomach, and often the retractor, stomodeum and esophagus (Figure 3 C; Table 2).

Unlike shell defects reported above, these animals were completely lacking the shell; however, they sometimes appeared to have a thin membrane where the shell would be. The periostracal membrane is a chitinous sheet that forms the outermost membrane of the shell, and is secreted by the periostracal groove, a fold in the anterior shell epithelium which can be stained with peroxidase substrates (Johnson et al., 2019). In wild-type larvae, strong staining is observed in the periostracal groove, which extends from left to right across the shell epithelium (Figure 3C[ii]). Weaker staining can be observed in the membrane itself (not shown). In UVI-3003-treated embryos, the periostracal groove was much shorter and only on the left side, and the periostracal membrane was reduced to a small patch on the left, posterior to the fragment of the groove (Figure 3C[iv]). These results show that UVI-3003 treatment disrupts the development of the periostracal groove, potentially explaining the impaired *shell* development. This phenotype could be a result of blocking RA binding to RXR; it could also reflect requirements for other RXR ligands during shell development.

To assess the requirements for *ToRXR* itself, we knocked down *ToRXR* using a translation-blocking morpholino (*ToRXR* MO1). Zygotes injected with *ToRXR* MO developed normally until shortly after the start of organogenesis and shell development (∼ 3.5-days after egg lay). At this point, they stopped progressing morphologically, remained in their arrested morphology for a few days and then dissociated (Figure 3D[iii]). We note that this is shortly after the start of the organogenesis window of RA sensitivity we described above. To confirm the specificity of the *ToRXR* MO1, we knocked down *ToRXR* using a second non-overlapping morpholino (*ToRXR* MO2).

Embryos injected with *ToRXR* MO2 exhibited a similar arrested phenotype to embryos injected with *ToRXR* MO1 (Figure 3D[iv]). These results indicate that RXR activation is necessary for shell development, and overall RXR function is necessary for progression into organogenesis. The more severe phenotype of RXR knockdown compared to the inhibitor could reflect functions with other co-receptors or ligands as well as the combined block of positive and negative transcriptional regulation.

### ToRAR morpholino knockdown

Several studies have shown that, in molluscs, *RAR* has lost its ability to bind RA (Andre et al., 2019; Gutierrez-Mazariegos et al., 2014a; Urushitani et al., 2013). RAR is strongly conserved within molluscs, indicating that it still has an important function.

Embryos injected with 0.25 mM *ToRAR* MO developed normally and grew to be wild-type (Figure 3E[i]), while embryos injected with 0.5 mM *ToRAR* MO developed for two days, until they arrested at the end of gastrulation (Figure 3E[ii]) and fell apart a few days later. These embryos arrested at the same timepoint that we found the “cytoplasmic tail” phenotype with ATRA, and well before the other window of sensitivity (ca. 3 days AEL). This result supports an essential role for *ToRAR,* but indicates that it is distinct from other RA pathway functions.

### The response of the shell epithelium to RA perturbation

Defects in shell development were the strongest and most consistent effects of our various manipulations of putative RA pathway components. To begin to assess whether these defects might have similar causes, we examined the effects of exogenous RA and CYP26A knockdown on the expression of two genes that are specifically expressed in the developing shell. In a wild-type embryo, *ToSushi domain-containing* is expressed in the entire mantle epithelium growth zone (MEGZ), most strongly at the anterior of its expression domain, and grading to lower expression posteriorly (Fig. 4A and Dao et al., 2025). Both 9-*cis* RA treatment and *ToCYP26A* knockdown reduced the area of expression of *ToSushi domain-containing,* and seemed to abolish the gradient observed in wildtype. *ToREV-ERB* is expressed in the anterior-most row of the epithelium (the anterior growth zone, AGZ, Dao et al, 2025). After 9-cis exposure or *ToCYP26A* MO knockdown, the single row of expression changes to a much more disorganized pattern, with no discrete row, and staining cells scattered around the mantle epithelium. (Figure 4A). These results indicate that these two manipulations have similar effects on these shell epithelium genes, further supporting the idea that ToCYP26A is normally involved in RA signaling in this embryo. In addition, these results indicate that disruption of shell development is not simply a loss or reduction of the shell secreting mantle — in one case the zone of expression is compressed and in the other it is disorganized and expanded.

**Figure 4.**
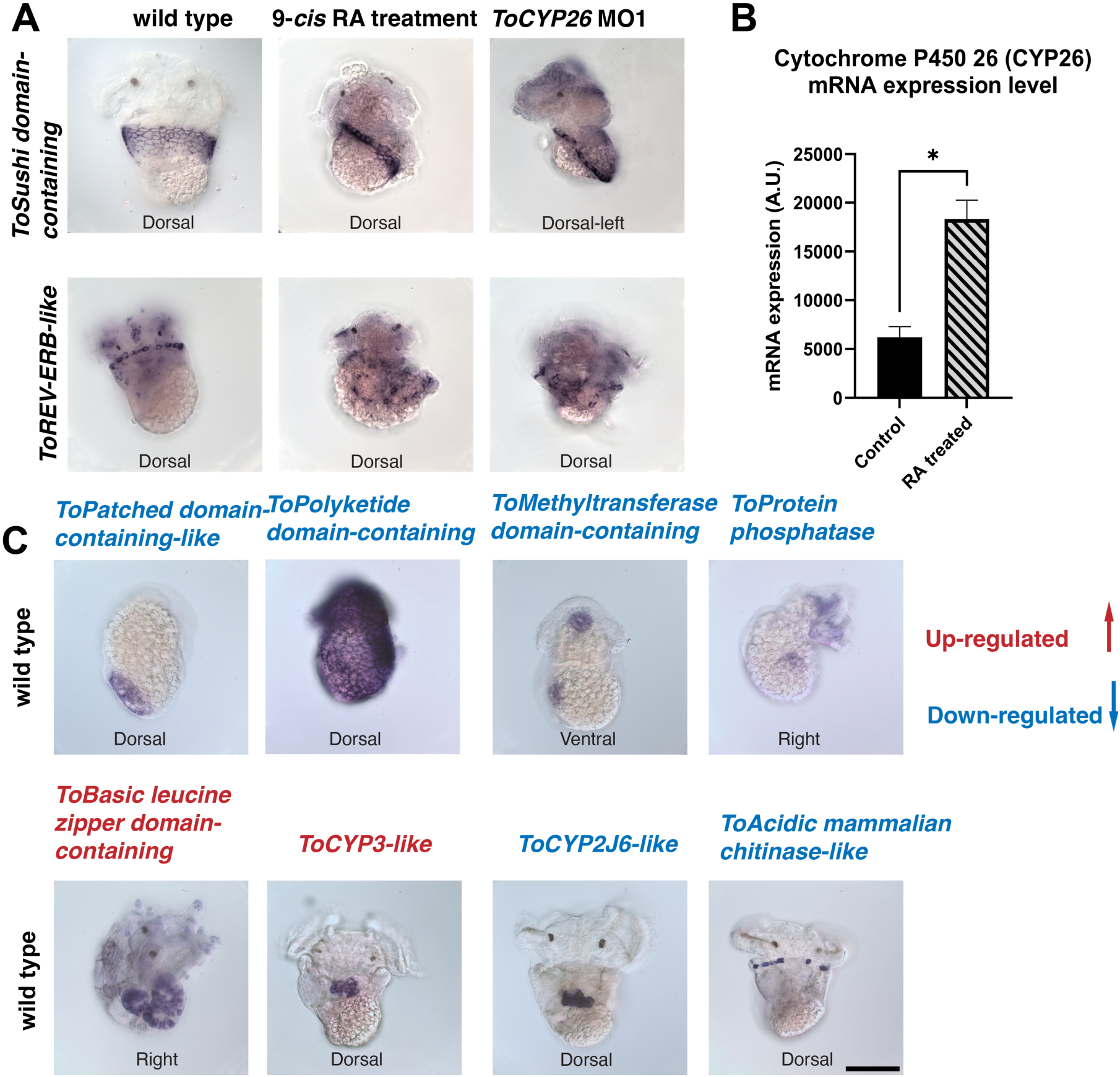
Gene regulation downstream of RA treatment in *Tritia*. (A) *ToCYP26A* mRNA was up-regulated after 2 hours of treatment with 9-*cis* RA. (B) Changes in expression patterns of *ToSushi domain-containing* and *ToREV-ERB-like* following 9-*cis* RA treatment or *ToCYP26A* MO1 knockdown. (C) Expression patterns of up-regulated (red) and down-regulated (blue) genes after RA treatment. *ToPatched domain-containing-like* (down-regulated at 12 hours) was expressed in the shell field of wild-type 4-day old organogenesis embryos. *ToPolyketide domain-containing* (down at 4h) was expressed ubiquitously but slightly darker near the shell shield in the late organogenesis stage. *ToMethyltransferase domain-containing* (down at 12h) was expressed in the stomodeum (mouth) and a small area on the right side of an embryo at the late organogenesis. *ToProtein phosphatase* (down at 4h) was expressed on the head and a small area on the right side of an embryo at the late organogenesis. *ToBasic leucine zipper domain-containing* (up at 2h) was in the digestive gland. *ToCYP3-like* (up at 8h) was expressed in a cluster of cells in the mantle cavity on the dorsal midline. *ToCYP2j6-like* (down at 12h) was expressed in a cluster of cells in the mantle cavity on the dorsal midline. *ToAcidic mammalian chitinase-like* (down at 12h) was expressed on the shell epithelium. Scale bar: 100 μm

### Transcriptional response to retinoic acid

In vertebrates, retinoic acid regulates many embryonic and physiological events by directly interacting with DNA and regulating downstream genes. Unliganded (apo) *RAR-RXR* heterodimer binds to repressors and represses gene transcription (le Maire et al., 2010). The *RAR-RXR* heterodimer recruits co-activators upon ligand binding (Cordeiro et al., 2019; Perissi and Rosenfeld, 2005). To detect early transcriptional change during retinoic acid treatment, we treated embryos for 2, 4, 8 and 12 hours with 9-*cis* RA and collected RNAs for RNA sequencing. Consistent with unliganded receptor generally acting as a repressor, we found that more genes were up-regulated after treatment than downregulated: at 2 hours there were 27 upregulated vs 8 downregulated; at 4 hours there were 74 up vs. 37 down. This trend continued at later stages, but it is likely that some of these interactions were not directly mediated by RA binding receptors. One well-known feedback loop in vertebrates is that when RA level is artificially elevated, the animals up-regulate their CYP26A to counter the heightened RA level. In vertebrates, this is mediated directly by RAR/RXR heterodimers binding to regulatory sequences of the CYP26A1 gene (Wang et al., 2002). In our experiment, *ToCYP26A* mRNA was up-regulated at 2 hours after retinoic acid treatment (Figure 4B). We also found a distinct gene that is most similar to CYP26B1 genes, and also upregulated. This latter transcript is ubiquitous, and has much lower read counts in controls than ToCYP26A (data not shown).

To further examine the transcriptional response, we chose differentially regulated genes with predicted functions indicating they could be developmental regulators, and performed *in situ* hybridization to determine their expression patterns. We discovered RA downstream genes that were expressed near the shell field, on the shell epithelium, in the digestive gland, and in structures in the dorsal mantle cavity. One notable gene we recovered was *ToPatched domain-containing-like*. We previously found that this transcript was lower after 2d micromere ablation, which prevents shell formation in *Tritia* (Dao et al., 2025). Here, expression level of *ToPatched domain-containing-like* declined at 12 hours after 9-*cis* RA treatment. *ToPatched domain-containing-like* is specific to the shell field in veligers (Dao et al., 2025) and organogenesis (Figure 4C). This result further strengthens the connection between RA signaling and shell development. Two other genes from this screen were also enriched in the developing shell. *ToPolyketide domain-containing* was expressed ubiquitously but slightly enriched at the shell margin in the late organogenesis stage. *ToAcidic mammalian chitinase-like* was expressed in particular cells in the anterior-most row (AGZ) in the shell epithelium in early veligers (Figure 4C). As described above, our functional studies have shown that both increasing and decreasing RA levels impacts shell development (and other phenotypes). In that light, it is striking that three of the genes we found with specific patterns in this RA response screen were expressed in the shell, and all were down-regulated after RA treatment. This indicates that the developing shell is an important target of RA and suggests that the pathway may be in part negative or modulatory for shell development.

Another transcript, *ToBasic leucine zipper domain-containing,* was in the digestive glands (Fig. 4C), where ALDH1A2 is also expressed. *ToMethyltransferase domain-containing* was expressed in the stomodeum (mouth) on the ventral side and a small area on the right side of a *Tritia* embryo during late organogenesis embryo. *ToProtein phosphatase-like* was expressed more broadly in the head and in a small area on the right side of a late organogenesis embryo. Two transcripts encoding cytochrome p450 proteins, *ToCYP3-like, and ToCYP2J6-like* were both expressed in groups of cells in the dorsal mantle cavity, in the same region as *ToRDH10* and *ToApolipoD-like*.

### Strong similarity of shell phenotypes and RA pathway phenotypes

Overall, the RA treatments look remarkably similar to each other, and different from all the other phenotypes we have characterized in other studies, except for those that target the shell. Together, this is strong support that these RA components are working together in a pathway, and one that targets the shell. To more explicitly test this, we performed a hierarchical clustering analysis, using the phenotypes from this paper, and the other comparable published phenotypes from our lab (i.e. those where we scored the same set of structures; Figure 5). All of the RA perturbations formed a large cluster, together with the treatments which were dominated by shell defects. The shell defect experiments were 2d deletion, which generally prevents any external shell formation, and *ToChitin binding domain* MO knockdown, which strongly impairs shell development. Both have other phenotypes secondary to their shell defects: they generally lack normal digestive glands and retractor muscles and often have defects in the intestine, esophagus, stomodeum and stomach (Dao et al., 2025). In the cluster, shell defect treatments were interspersed within the RA treatments. The remaining treatments are outgroups to the RA/shell cluster, demonstrating that RA perturbation phenotypes are distinct and have only been seen after RA or shell perturbations. We note that within the RA treatments, there was no clustering of treatments that would be predicted to raise RA vs. those that would be predicted to lower it.

**Figure 5.**
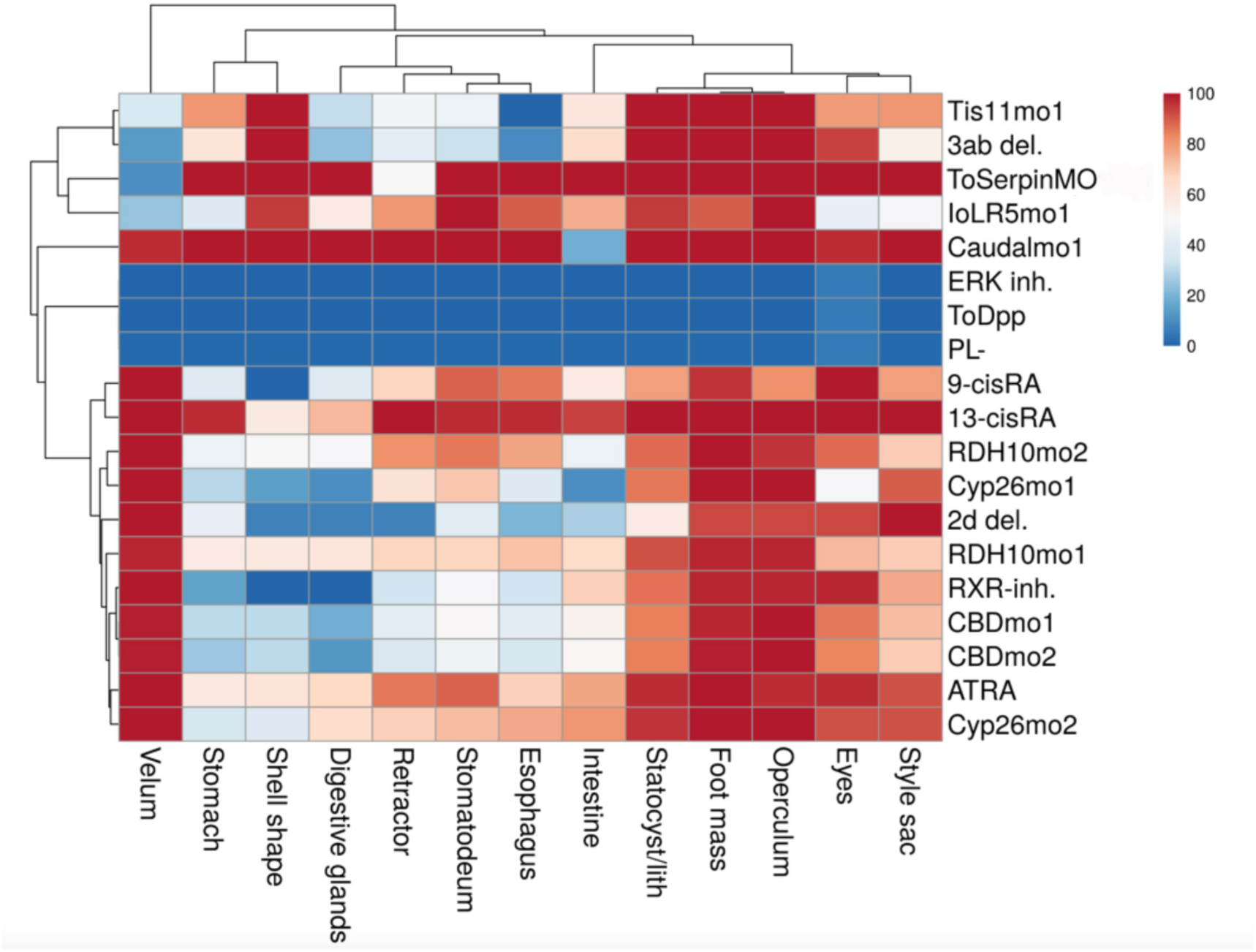
Hierarchical clustering of phenotypes Clustering of organ scoring from multiple RA signaling manipulations (RA treatment, *ToCYP26A* knockdowns, *ToRDH10* MO knockdowns, and RXR antagonist treatment) with shell manipulations (2d ablation and *ToChitinBD* MO knockdowns (Dao et al., 2025)) and other non-shell manipulations ToSerpinMO, 3ab ablation, *ToTis11* MO knockdown, *ToLR5* MO knockdown, *ToCaudal* MO knockdown, polar lobe ablation, ERK inhibition, ToDppMO knockdown (references in Table 2 notes).

## Discussion

Along with RXR and RAR, the Cyp26 and RDH10 genes form a core set of putative RA-signaling genes that are found across non-chordates (Albalat, 2009). We have performed the first knockdowns of CYP26 and RDH10 in a non-chordate and compared them to knockdowns of RXR, RAR, RA treatment and RXR inhibition in the same system. The concordance of the phenotypes of RA treatments, *ToCYP26A* knockdown, *ToRDH10* knockdown and RXR inhibition provide the strongest functional evidence to date of an extensive set of RA signaling genes functioning together outside chordates. This is further strengthened by the similarities in gene expression patterns of the putative RA pathway genes and RA response genes. Multiple genes are specific to the developing shell, consistent with an important role for RA here, and several are expressed in the dorsal mantle cavity, suggesting that this is a focus of RA metabolism and signaling.

The role of RAR in molluscs has been mysterious since it was discovered that all mollusc RARs examined have lost the ability to bind RA. The knockdown of *ToRAR* we report here is the first in a mollusc embryo. Most notable are the differences from our other RA perturbations: the phenotype — embryonic arrest — is different from all but the RXR knockdown, and the timing — end of gastrulation — is earlier than the RXR MO arrest phenotype in organogenesis, and the shell development window of sensitivity.

These observations are consistent with a role in a process that is distinct from RA signaling. Unfortunately, beyond the timing, the arrest phenotype does not provide an indication of what this process might be. Since the gastrulation window of ATRA sensitivity coincides with the timing of RAR knockdown arrest, it is tempting to think that they may be related. However, since *ToRAR* almost certainly doesn’t bind RA, it is hard to imagine a simple way they could be linked.

The fact that the *ToRXR*MO arrest was later than the *ToRAR*MO arrest suggests that ToRXR does not participate in whatever essential process is arrested in ToRAR embryos. However, it is possible that ToRAR is participating in later events with ToRXR, perhaps as a heterodimer. Mollusc RAR/RXR heterodimers can bind DNA at canonical RA Response Elements (RAREs); mollusc RXR/RXR homodimers can also bind DNA at RAREs (Gutierrez-Mazariegos et al., 2014a). Thus, the role of RA signaling in shell development could be mediated by RXR/RXR or RAR/RXR dimers. The ToRXRMO arrest was likely a consequence of disrupting a non-RA related role of RXR, since it was so much more severe than the other RA perturbations.

The annelid *Platynereis* is the only other spiralian where RXR and RAR have been knocked down by specific gene knockdown (Handberg-Thorsager et al., 2018). Both knockdowns resulted in defects in neurogenesis, both in axon arrangement and number of neurons. It is possible that there are neural defects after our RA perturbations in *Tritia*, but we have not examined neural patterning because the morphology of the posterior of the body is so abnormal that it would not be possible to exclude indirect effects on the nervous system.

The speed of the appearance of cytoplasmic tails after ATRA treatment at gastrulation (∼15 minutes) precludes a transcriptional response. There is evidence for non-transcriptional actions of RA. Some effects of RA on mollusc neurons and growth cones seem to be too rapid to be transcriptionally mediated (Farrar et al., 2009).

Recent studies have indicated the presence of RARα in membrane lipid rafts of some cell lines. This pool of extra-nuclear RAR activated p38MAPK in response to all-*trans* RA treatment. Interestingly, the activation of p38MAPK occurred within only minutes after the addition of all-*trans* RA (Alsayed et al., 2001; Piskunov and Rochette-Egly, 2012). Perhaps a similar mechanism might operate through RXR in molluscs, where RAR does not bind RA.

In vertebrates, the retinoic acid-degrading enzyme, *CYP26A*, is positively regulated by RA levels, so an increase in RA level induces *CYP26A* upregulation to counter and bring down RA levels. In *Tritia,* we detected an increase in expression level of *ToCYP26A* mRNA following RA treatment, indicating similar regulation. We note that we observed the crooked shell phenotype in RA-treated embryos and RDH10 MO embryos but not in *ToCYP26A* knockdown embryos. The crooked shell phenotype, with its irregular biomineral ridges across the middle of the shell, is likely caused by a temporary disruption of shell development followed by recovery and resumption of normal development. One possible mechanism for this is that RA treatment disrupted shell development temporarily, but subsequent *ToCYP26A* upregulation brought levels down and allowed normal shell development to resume.

The spatial relationships between putative RA pathway components could provide hints about their patterning roles. In vertebrates, retinoic acid often acts in gradients created by non-overlapping distribution of RA-synthesis and RA-degrading enzymes (as reviewed in Cunningham and Duester, 2015; Sakai et al., 2001). In *Tritia,* the expression patterns of the main RA pathway components did not clearly indicate a long range anterior-posterior gradient. However, *ToCYP26A*, an RA-degrading enzyme and *ToBCMO1-like*, an RA-synthesizing enzyme were both expressed in similar patterns near the edge of shell field in the organogenesis stage; they may be offset a small amount to create a very localized gradient. The expression of *ToApolipoD-like* (RBP4-related) at the shell margin also supports the idea that RA levels are modulated there. Many genes in the shell epithelium are expressed in gradients (Dao et al, 2025). The compression of the *ToSushi pattern* in both RA treatments is consistent with the loss of a gradient in the shell epithelium. The change in the *ToRev-ERB* is not consistent with this however — if the *ToSushi* pattern is anteriorized, the *ToRev-ERB* pattern is posteriorized. Nevertheless, the shell epithelium is complex and different genes and populations of cells could be regulated by distinct mechanisms (Dao et al., 2025; Jackson et al., 2007; Johnson et al., 2019).

Perturbations that would raise RA levels (RA treatment, *ToCYP26A* MO) and perturbations that would lower levels (*ToRDH10* MO, RXR inhibitor) had indistinguishable effects on shell development. This suggests that the RA level in*Tritia* must be maintained at a specific level, possibly as a permissive cue. This could be consistent with a fine-scale gradient.

If *ToRDH10* is an important factor for RA synthesis, it seems likely that there are additional RALDH factors that would be more broadly expressed than the one that we have reported here, e.g. in the developing shell. In addition, Cytochrome P4502J genes have been proposed to participate in RALDH1A-independent oxidation of retinaldehyde in vertebrates (Albalat, 2009), and *ToCYP2J6* is expressed in a very similar location to *ToRDH10* in the dorsal mantle cavity, under the developing shell, making this a potential source for RA. *ToFatty Acid Binding protein-like* (CRABP1-related) is also expressed in a similar pattern.

The expression domains of *ToRDH10, ToFatty Acid Binding protein-like* and *ToCYP2J6* in the dorsal mantle cavity are positioned at the interface between the yolk mass and the rest of the embryo during the organogenesis stages when the embryo is sensitive to RA. This suggests that these cells may be involved in mobilizing retinoid stores from the yolk during embryogenesis. These cells are also very near the growing edge of the shell epithelium during late organogenesis. Making the shell margin an RA signaling center for the whole embryo could be an organizing principle for the gastropod embryo. Keeping the source of this signal in the same place relative to other organs as the animal develops and elongates towards the anterior could provide relative spatial cues similar to posterior growth zone RA signaling in vertebrates. In both cases, the fixed location of a signaling center as the animal elongates allows growth of different regions and organs to be coordinated.

## MATERIALS AND METHODS

### In situ *probe synthesis*

We extracted total RNA from embryos at various stages using Trizol (Invitrogen) and reverse transcribed it into cDNA using Superscript^TM^ IV Reverse Transcriptase kit (ThermoFisher). Primers for probe template amplification were designed using Batchprimer3 (https://probes.pw.usda.gov/batchprimer3/) to be produce templates 700-100 bp long, and including the T7 promoter sequence (TAATACGACTCACTATAGGG) flanking the 5’ end of the forward primer and the T3 promoter sequence (AATTAACCCTCACTAAAGGG) flanking the 5’ end of the reverse primer. We performed in vitro RNA *in situ* probe synthesis with the PCR products using a mixture of NTPs containing digUTP (Roche).

### In situ hybridization

*In situ* hybridization was carried out as described (Lambert and Nagy, 2002). Briefly, fixed embryos stored in methanol were rehydrated, then treated with TEA (0.1 M triethanolamine hydrochloride pH 8.0) and acetic anhydride to suppress non-specific probe binding. Embryos were washed with PBTw and then incubated with DIG-conjugated probes diluted in Hyb buffer (50% Formamide, 5x Denhardt’s, 5x SSC, 0.1% Tween, H_2_O) with blockers (200 μg/ml tRNA, 100 μg/ml Heparin, 200μg/ml Salmon sperm) at 68°C in 72 hours. Embryos were blocked with 2% BSA in PBTw and incubated with anti-DIG-AP antibody (Roche) diluted at 1:3000 ratio in 2% BSA in PBTw overnight at room temperature. Embryos were washed with AP developing buffer (100 mM NaCl, 50 mM MgC_l2_, 100 mM Tris pH 9.5 and 0.5% Tween), and developed with AP developing buffer plus 1 mg/ml BCIP and NBT substrates (Roche).

### Retinoic acid treatment

All handling of RA isomers was performed under yellow light, including dilution of stock solutions in DMSO, dilution of DMSO stocks into filtered artificial sea water, and addition of embryos to RA-sea water. When not being handled, embryos were kept in the dark during RA treatment. We collected embryos at 1 through 4-cell stages and reared them to the indicated stages to start treatments. For multi-day treatments, embryos were moved to fresh RA-sea water mixtures each day.

### Morpholino injection

Morpholino oligos were designed by the vendor (Gene Tools). Zygotic injection was performed as described previously (Rabinowitz et al., 2008) except that injections were performed in un-supplemented filtered artificial sea water. Injected animals were reared until they depleted their yolk for scoring. The indicated concentrations for injected solutions are the concentrations in the pipette.

### RA treatment RNA-seq

We divided each capsule equally into 24 dishes (12 controls and 12 experiments). We started embryo treatments at the same time and stopped treatment at 2 hrs, 4 hrs, 8 hrs and 12 hrs. Each timepoint had 3 replicates. Total RNA was extracted using TRIzol (Invitrogen). RNA concentration was assessed using a NanoDrop ND-1000 spectrophotometer, an Agilent 2100 Bioanalyzer and a Qubit flourometer. RNA was poly-A selected and sequenced with the Illumina HiSeq 2x150 bp protocol. The cleaned fastq files can be accessed with the accession number PRJNA1088840 on https://www.ncbi.nlm.nih.gov/sra (https://www.ncbi.nlm.nih.gov/sra/PRJNA1088840). RNA-seq analysis was performed as described (Dao et al., 2025).

## Notes

### Competing Interest Statement

The authors have declared no competing interest.

### Funding

- U.S. National Science Foundation: IOS 2053371

